# Lipid Droplets mediated Japanese Encephalitis Virus small Extracellular Vesicle Release

**DOI:** 10.1101/2025.11.04.686473

**Authors:** Bhaghyasree Mallick, Ankita Sarkar, Ananya Mondal, Tamoghna Chakraborty, Khadijah Khan, Dilip Kumar, Subhas Chandra Biswas, Sourish Ghosh

**Affiliations:** Infectious Diseases & Immunology Division, CSIR-Indian Institute of Chemical Biology, 4, Raja S.C. Mullick Road, Jadavpur, Kolkata-700032, West Bengal, India; Cell Biology & Physiology Division, CSIR-Indian Institute of Chemical Biology, 4, Raja S.C. Mullick Road, Jadavpur, Kolkata-700032, West Bengal, India; Trivedi School of Biosciences, Ashoka University, Sonipat, Haryana; Academy of Scientific and Innovative Research (AcSIR), Ghaziabad 201002, India

**Keywords:** Small extracellular vesicles, lipid droplets, intracellular trafficking, multivesicular bodies, Japanese Encephalitis Virus, nSMase2, Neurons

## Abstract

Lipid droplets (LDs) and small Extracellular Vesicles (sEVs) are classically known for lipid metabolism and intercellular communication, respectively. Here, we reveal a mechanistic connection between LD dynamics and sEV-mediated non-lytic release of Japanese Encephalitis Virus (JEV) from neuronal cells. Using Neuro2A, SH-SY5Y, N9 microglia, and primary cortical neurons, we show that JEV is packaged within sEVs (∼200 nm) through an ESCRT-independent, neutral sphingomyelinase 2 (nSMase2)/ ceramide-dependent pathway. Virions inside sEVs display a higher JEV Premature Membrane/Membrane protein (PrM/M) ratio compared to those released via the conventional secretory pathway. Although containing a higher proportion of premature virions than mature ones, sEV-associated JEV virions gain an evolutionary advantage by evading immune detection and delivering multiple virions to recipient cells, thereby increasing overall infection efficiency. Temporal profiling showed early cytoplasmic LD enrichment (from 6 hpi), followed by a surge in sEV release from 14 hpi, suggesting sequential roles for LDs and sEVs. nSMase2 inhibition decreased sEV-mediated egress without affecting viral replication, but increased cytoplasmic LD abundance, consistent with LD underutilization in multivesicular bodies (MVB) biogenesis. Our findings identify LDs as facilitators of MVB formation and nSMase2 as a key driver of sEV-mediated viral exit, revealing parallel yet coordinated pathways in JEV’s stealthy egress.

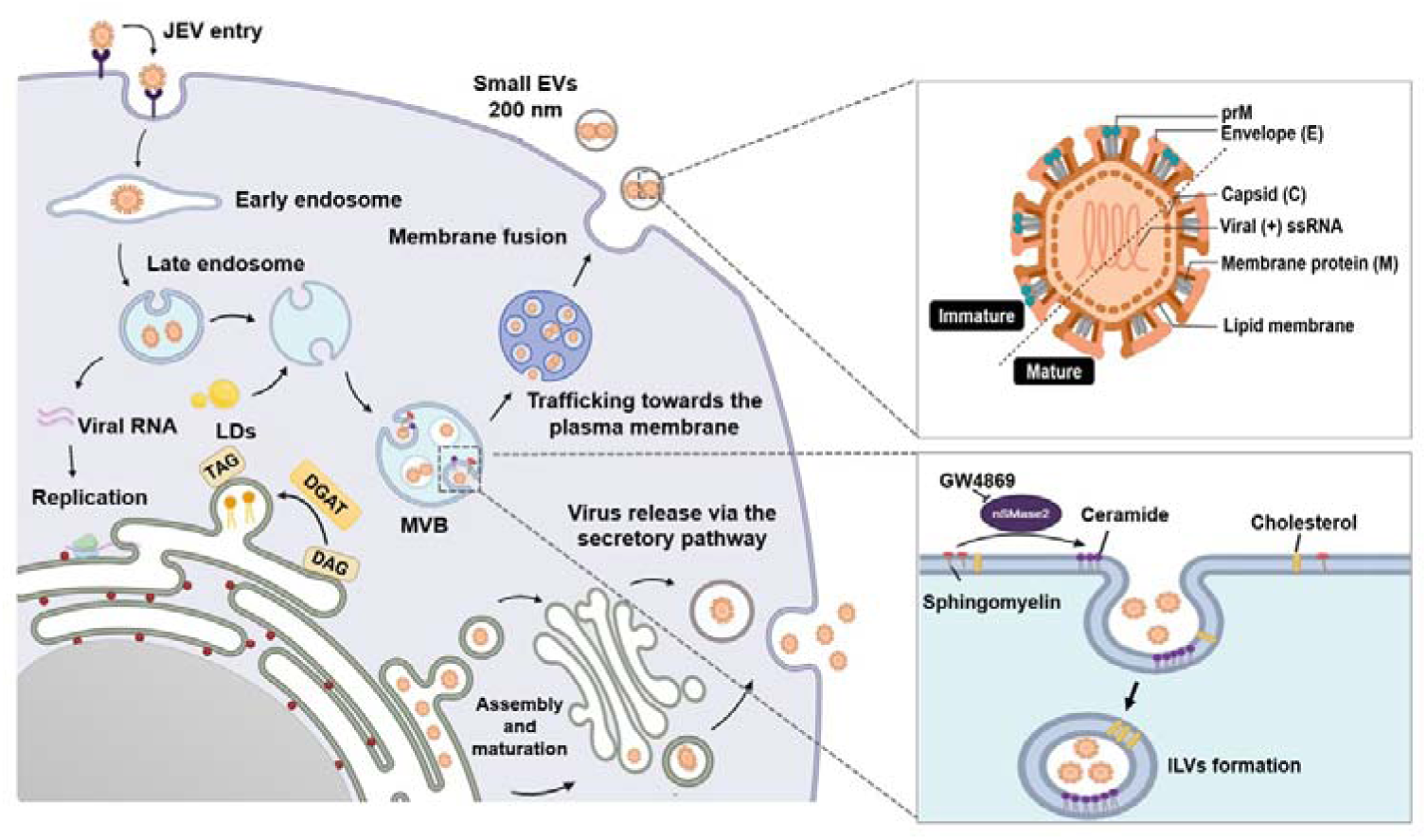

## 1. Introduction

Host-pathogen interaction is influenced not only by the immune system but also by a tightly coordinated network of membrane-associated organelles that control cargo storage, trafficking, and metabolic balance (Stanton, 2021). Among the most intriguing of these are small extracellular vesicles (sEVs) —once considered trivial cellular by-products, now conceded as powerful modulators of intercellular communication, metabolic signalling, and viral trafficking (Wessler & Meisner-Kober, 2025; Hu *et al*, 2023; Xia *et al*, 2023; Couch *et al*, 2021). Extracellular vesicles (EVs), including exosomes and microvesicles, are nanoscale, lipid bilayer-enclosed carriers secreted by nearly all cell types with particle sizes typically ranging from 30 to 1000 nm (Raposo & Stoorvogel, 2013). Generated either by ESCRT-dependent machinery or -independent pathways, EVs transport bioactive molecules such as proteins, RNA, and lipids across cells and tissues, shaping immune responses, tissue remodelling, and—most pertinently—viral spread (György *et al*, 2011; Altan-Bonnet *et al*, 2019; Santiana *et al*, 2018). sEVs have emerged as important mediators of intercellular communication and have been implicated in the spread of infection in several flaviviral models, such as Dengue virus (DENV), Zika virus (ZIKV), West Nile virus (WNV), including Japanese Encephalitis Virus (JEV) (Slonchak *et al*, 2019; York *et al*, 2021; Calderón-Peláez *et al*, 2024; Xiong *et al*, 2025; Latanova *et al*, 2024; Martínez-Rojas *et al*, 2025). Neutral sphingomyelinase 2 (nSMase2), also known as SMPD3, has been shown to regulate the biogenesis and release of sEVs in neurons (Tallon *et al*, 2021b, 2021a). Inhibition of SMPD3, either through siRNA-mediated knockdown or pharmacological blockade using GW4869, resulted in reduced ZIKV loads in neurons and neuron-derived extracellular vesicles (sEVs) (Zhou *et al*, 2019). A similar observation was made in the case of JEV, where GW4869 inhibited both replication and virus release inside sEVs in various cell types, emphasizing role of the nSMase2-ceramide pathway in EV-mediated flaviviral transmission (Xiong *et al*, 2025).

In contrast to extracellular vesicles, lipid droplets (LDs) are intracellular organelles that serve as energy reservoirs, consisting primarily of neutral lipids such as triacylglycerols (TAGs) and cholesteryl esters, encased within a phospholipid monolayer (Olzmann & Carvalho, 2019). LDs have evolved from mere lipid stores into dynamic, multifunctional platforms that play a role in protein sequestration, immune signaling, lipid trafficking, and viral replication, including flaviviruses (Monson *et al*, 2021; Peng *et al*, 2025; Martins *et al*, 2024; Carvalho *et al*, 2012; Sarkar *et al*, 2021). Pharmacological inhibition of DGAT-1, an enzyme involved in LD biogenesis, has been shown to significantly reduce LD formation and ZIKV replication in both *in vitro* and *in vivo* systems (Schöbel *et al*, 2024). Beyond their role in viral replication, LDs also contribute to the early antiviral immune response, particularly through their influence on type I interferon (IFN) signalling following infection, thereby helping to control viral spread (Monson *et al*, 2021). Consistent with the immunomodulatory functions of LDs, blockade of LD formation also profoundly affects inflammatory cytokine production in the brain, highlighting the dual role of LDs in both viral pathogenesis and host immune regulation (Tan *et al*, 2024).

While the individual roles of EVs and LDs have been extensively studied in the context of viral infections, their interconnection has not been thoroughly explored. A recent study revealed that an abundance of LDs strongly correlates with the release of sEVs in various cancer cell lines (Genard *et al*, 2024). Pharmacological inhibition of LD metabolism significantly suppressed sEV secretion, while metabolic stressors such as hypoxia, acidosis, or irradiation—known to enhance LD biogenesis— dramatically elevated vesicle release. These findings suggest that lipid metabolic flux may govern the dynamics of EV biogenesis. Further linking these organelles is the ceramide pathway, a shared molecular axis involving neutral sphingomyelinase 2 (nSMase2), a key enzyme that converts sphingomyelin into ceramide—a lipid critical for both EV budding and LD turnover (Huang *et al*, 2025; Schempp *et al*, 2024). Notably, nSMase2 is highly expressed in the central nervous system (CNS) and plays essential roles in brain development, membrane remodelling, and neuronal communication. Viruses, including flaviviruses, are known to hijack the ESCRT-independent, ceramide-dependent route to enhance their transmission (Beckmann & Becker, 2021).

Japanese Encephalitis Virus (JEV) is a mosquito-borne, neurotropic flavivirus and the leading cause of viral encephalitis in Asia and the Western Pacific (Mulvey *et al*, 2021). Globally, more than 68,000 cases of Japanese Encephalitis (JE) are reported annually, with a case fatality rate approaching 30%. Among survivors, 20–30% suffer long-term neurological sequelae, including motor impairments, recurrent seizures, and cognitive or speech deficits (Moore, 2021). Although significant progress has been made in understanding JEV pathogenesis and host-pathogen interactions, the mechanisms underlying viral egress, particularly from neuronal cells, remain poorly defined. Recent studies suggest that JEV can exit host cells encapsulated sEVs, but the molecular determinants of this process remain unclear. Given the emerging recognition of LDs and EVs as central hubs in lipid metabolism and intercellular signalling, we sought to determine whether JEV exploits LD–sEV axis to facilitate its release from neuronal and glial cells, with a specific focus on the ESCRT-independent pathway mediated by ceramide-producing enzyme nSMase2.

## 2 Results

### 2.1. JEV egress from neuronal cells via a non-lytic mechanism

Japanese Encephalitis Virus (JEV), a well-known neurotropic virus, efficiently infects various neuronal cell lines, including Neuro2a, SHSY-5Y, and N9 microglial cells, as well as primary cortical neurons (Mohapatra *et al*, 2023; Kalia *et al*, 2013). These models were used in this study to examine the dynamics of JEV replication and release (Fig. 1A). To visualize JEV particles, viral stocks were pre-incubated with the lipophilic PKH 26 red dye, which incorporates into the host-derived viral envelope. After removing unbound dye, the labelled inoculum was added to Neuro2a cells. Discrete fluorescent puncta, representing individual or clustered virions, were observed as early as 6 hours post-inoculation (hpi), with a steady increase in both intensity and number at 18 and 24 hpi (Fig. S1A-B), indicating active viral replication. Additionally, we incubated the fixed cells expressing the PKH-stained puncta from 18 hpi with anti-JEV-Capsid and observed colocalization with the puncta, confirming that these puncta represent replicating viral populations (Fig. 1B).

**Figure 1:**
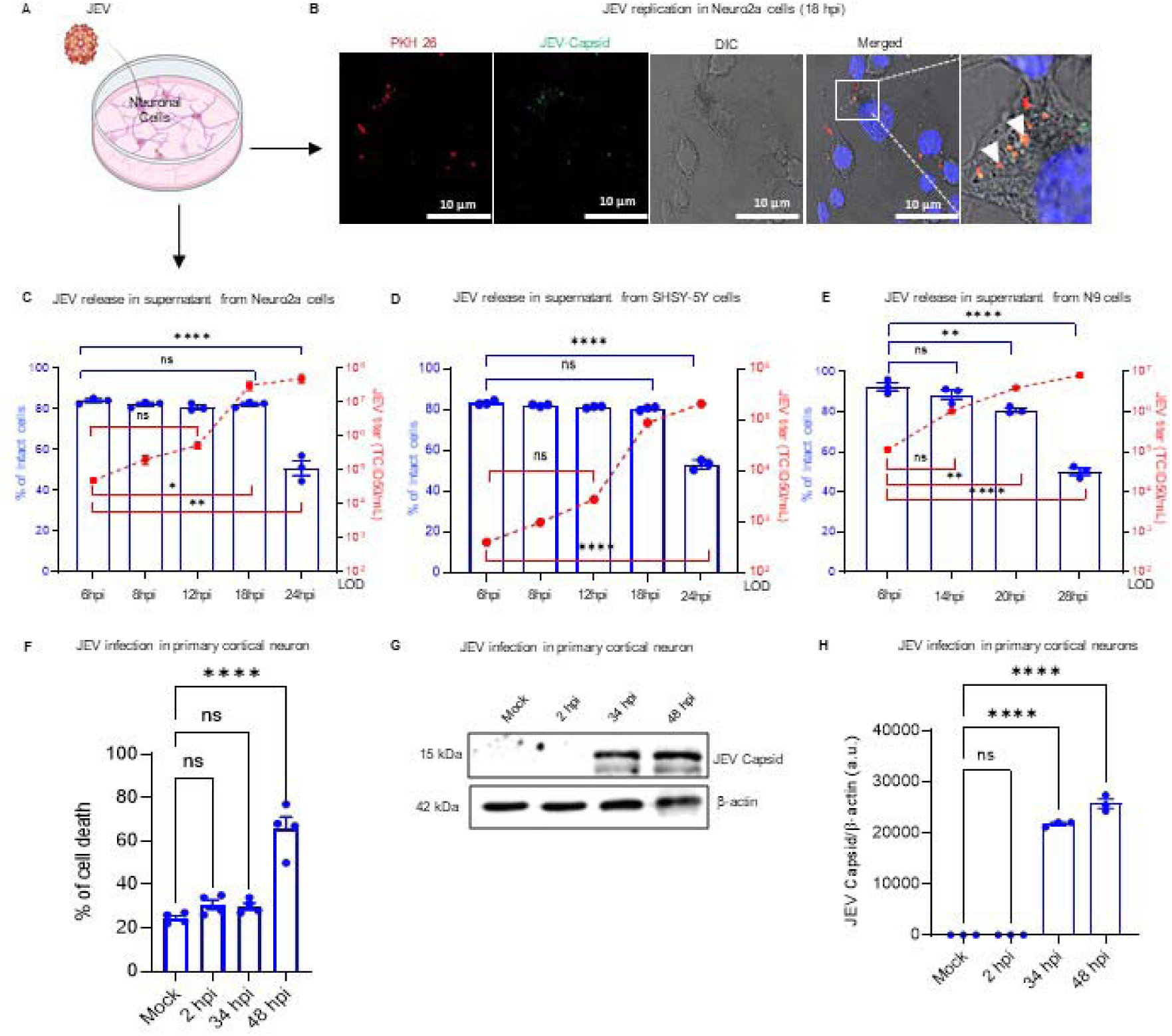
**JEV egress from neuronal cells via a non-lytic mechanism** (A) Neuronal cells were inoculated with JEV to determine optimal time points for egress analysis. (B) JEV stock (an enveloped virus) was pre-labeled with PKH 26 red dye, washed, and applied to cells for visualization and counting of virus puncta. Virus puncta were observed at 18 hours post-inoculation (hpi) and colocalized with JEV-Capsid protein. Scale Bar = 10µm. (C–E) Viral titers (TCID50/ mL) were measured in the culture supernatants collected at various time points from (C) Neuro2a, (D) SH-SY5Y, and (E) N9 microglial cell lines. Corresponding cell viability was assessed using Trypan Blue exclusion. LOD= Limit of Detection of TCID50/mL. (F) Primary cortical neurons remained morphologically intact up to 34 hpi following JEV inoculation. (G-H) Viral replication at 34 hpi was confirmed by immunoblot detection of JEV capsid protein and quantified by densitometry in (H); β-actin served as a loading control. Statistical significance was calculated using one-way ANOVA followed by multiple comparison tests. **pL<L0.01, ***pL<L0.001, ****pL<L0.0001, ns= not significant.

Measurement of viral titers in the supernatant of infected Neuro2a, SHSY-5Y, and N9 cells showed a significant increase at 18 hpi in Neuro2a and SHSY-5Y cells, and 20 hpi in N9 cells, respectively (Fig. 1C–E). Notably, cell viability remained mostly unaffected at these time points, as shown by the Trypan Blue exclusion assay, indicating that viral release occurred without substantial lysis. To confirm these findings in a more physiologically relevant system, primary cortical neurons were infected with JEV. These neurons stayed intact up to 34 hpi (Fig. 1F, Fig. S1C), while strong viral replication was visible by capsid protein expression at 34 and 48 hpi (Fig. 1G–H). To support these results, we blocked the conventional secretory pathway, which is mainly responsible for protein secretion and is known to release mature JEV virions. Specifically, we inhibited trafficking through the endoplasmic reticulum-to-Golgi compartment (ERGIC) in Neuro2a cells using 10μg/mL of Brefeldin A (BFA) (Fig. S1D-E), a reversible and potent inhibitor of ERGIC transport (Nebenführ *et al*, 2002). Our experiments showed that 6 hours of BFA treatment did not affect JEV replication but significantly reduced the release of infectious particles into the extracellular space (Fig. S1F-G). Despite this reduction, titers remained above the detection limit (LOD) for TCID50 analysis, indicating that JEV can still release infectious virus. These findings suggest that while the ER-Golgi pathway may facilitate viral release, JEV can also employ alternative routes, such as extracellular vesicle-mediated secretion.

### 2.2. JEV is released via small extracellular vesicles (sEVs)

To investigate whether JEV utilizes extracellular vesicles for its release, supernatants from cells were processed using a sequential centrifugation protocol as illustrated in Fig. 2A. The final 100,000 × g pellet was resuspended in serum-free media and subjected to purification using ExoQuick ULTRA columns, following the manufacturer’s instructions (see Methods). Vesicles isolated from JEV-infected Neuro2a (Fig. 2B), SHSY-5Y (Fig. 2C), primary cortical neurons (Fig. 2D), and N9 microglial cells (Fig. 2E) were analyzed for the presence of extracellular vesicle markers Alix and CD81, along with JEV capsid protein. Immunoblot analysis revealed a notable increase in all three markers in JEV-infected samples compared to mock controls (Fig. S2A–C), indicating enhanced release of virus-containing vesicles.

**Figure 2:**
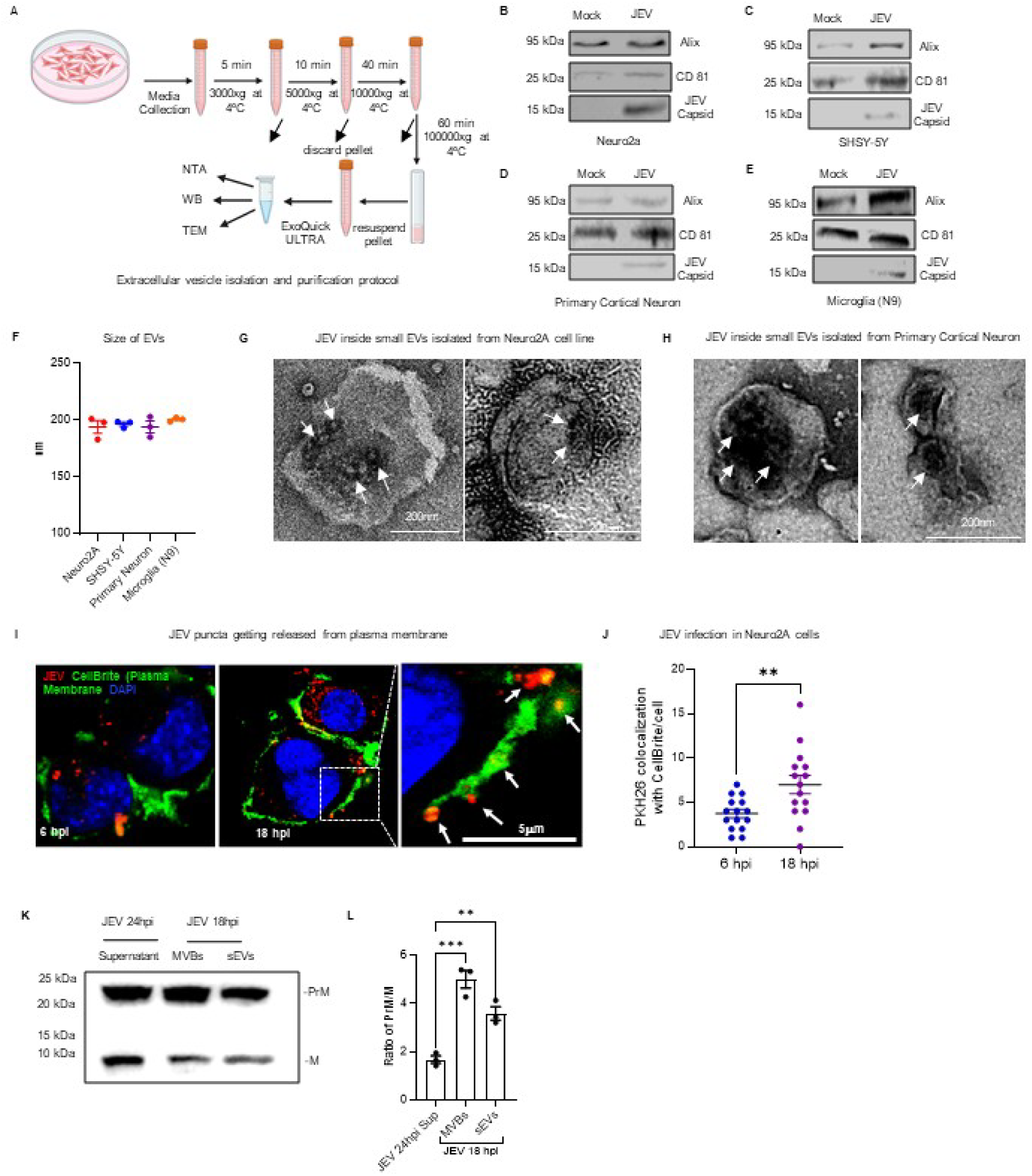
**JEV Is released via small extracellular vesicles (sEVs)** (A) Schematic representation of the protocol used to isolate and enrich small extracellular vesicles (sEVs) from cell culture supernatants. (B-E) sEVs were isolated from both mock- and JEV-infected (B) Neuro2a cells, (C) SH-SY5Y cells, (D) primary cortical neurons, and (E) N9 microglial cells, immunoblots showing expression of vesicle markers CD81 and Alix, along with JEV capsid expression from isolated sEVs. An equal volume of samples was added to wells for normalization. (F) The size distribution of vesicles was measured using nanoparticle tracking analysis (NTA). (G, H) Transmission electron microscopy (TEM) with negative staining visualized sEVs carrying JEV particles in (G) Neuro2a cells and (H) primary cortical neurons. Arrows indicate the presence of JEV particles inside the vesicles. Scale Bar = 200nm. (I, J) JEV particles, pre-labeled with PKH 26 red dye, were visualized using confocal microscopy at 6 and 18 hpi. (I) Representative confocal images show JEV puncta budding from the plasma membrane, stained with CellBrite dye, and (J) corresponding quantification of these events. Arrows indicate the JEV-puncta within or associated with membrane. Scale Bar = 5µm. (K, L) Measuring cleavage of the Premature Membrane protein (PrM) into the mature Membrane protein (M) of JEV in cell supernatant, multi-vesicular bodies (MVBs), and small extracellular vesicles (sEVs) collected at 24 hpi and 18 hpi, respectively. (K) shows the immunoblot expression and densitometric quantification in (L). An equal volume of samples was added to wells for normalization. Statistical significance was assessed using unpaired t-tests for pairwise comparisons and one-way ANOVA followed by multiple comparison test for multiple comparisons. **pL<L0.01, ***pL<L0.001, ****pL<L0.0001.

Nanoparticle tracking analysis (NTA) confirmed that the average size of vesicles from all cell types was around ∼200 nm (Fig. 2F; Fig. S2D–G), consistent with the size of small extracellular vesicles. Additionally, ultrastructural validation was performed using transmission electron microscopy (TEM) imaging with negative staining. Vesicles isolated from both Neuro2a (Fig. 2G) and primary cortical neurons (Fig. 2H) were observed to contain 2–3 JEV particles per vesicle, supporting JEV packaging within sEVs. To visualize the process of viral egress, confocal microscopy was performed using CellBrite (plasma membrane stain) and PKH 26-labeled JEV. At 18 hpi, but not at 6 hpi, distinct JEV puncta were seen pinching off from the plasma membrane and showed significant colocalization with CellBrite signal (Fig. 2I, J, Fig. S2H), indicating vesicular release of JEV particles from the cell surface. Consistent with the observation that JEV virions can be packaged into extracellular vesicles (EVs) and released through ESCRT-dependent or -independent pathways, a key question remains—why an enveloped virus like JEV use a host-derived vesicle to egress? Furin, a host enzyme, cleaves JEV Premature Membrane protein (PrM) to Membrane protein (M), resulting in mature viral particles, and the PrM/M ratio signifies the infectivity of JEV (Xiong *et al*, 2022)—the lesser the ratio, the more the infectivity. Our results show that the PrM/M ratio in JEV-infected cell supernatant at 24 hpi is lower in MVBs and sEVs collected at 18 hpi, indicating a higher proportion of immature viral particles being packaged inside MVBs and sEVs (Fig. 2K, L, Fig. S2I). These findings suggest that even immature virions may be transmitted via this mode, which may provide an adaptive advantage to the virus. Overall, these findings offer strong evidence that JEV is packaged and secreted within sEVs across various neuronal cell types. However, until now, it remains unclear whether the immature JEV virions packed within vesicles are capable of causing infection.

### 2.3. JEVs packaged within sEVs are infectious

We compared the relative infectivity of JEV released via sEVs versus free virus particles obtained from the sEVs. JEV-containing sEVs were isolated and divided into two fractions: one left intact and the other lysed using a detergent-free lysis buffer. To ensure that the lysis buffer did not compromise viral infectivity, we treated an equal amount of JEV stock with the same buffer and compared its replication capacity to untreated virus in Neuro2a cells. No significant difference in viral replication or titer was observed (data not shown), validating the use of the buffer for selective vesicle lysis.

Next, equal amounts of either intact sEV-associated JEV or vesicle-free JEV (lysed fraction) were inoculated onto Neuro2a cells (Fig. 3A). At 24 hpi, cells infected with JEV packaged in sEVs yielded significantly higher viral titers compared to those infected with the free virus (Fig. 3B), indicating superior infectivity of the vesicle-associated form. To further validate this *in vivo*, 4-day-old BALB/c mouse pups—an established model for JEV infection—were intracranially inoculated with either form of the virus (Fig. 3C). This route bypasses the blood-brain barrier and ensures equal delivery to the brain. At 7-day post-inoculation, brain tissues from the group inoculated with sEV-associated JEV showed markedly higher viral titers than those inoculated with free virus particles (Fig. 3D). Moreover, mice receiving the sEV-associated JEV exhibited more rapid and pronounced weight loss (Fig. 3E) and succumbed to infection earlier (Fig. 3F), consistent with a more severe disease phenotype. Together these results demonstrate that JEV enclosed within sEVs is significantly more infectious than its vesicle-free counterpart, both *in vitro* and *in vivo*, suggesting that sEV-mediated transmission of JEV even having a higher proportion of immature virions to mature virions are infectious. M protein, which mainly helps in JEV attachment to host cell membrane proteins for entry, vesicle-cloaked virions can bypass the requirement by getting packaged inside sEVs.

**Figure 3:**
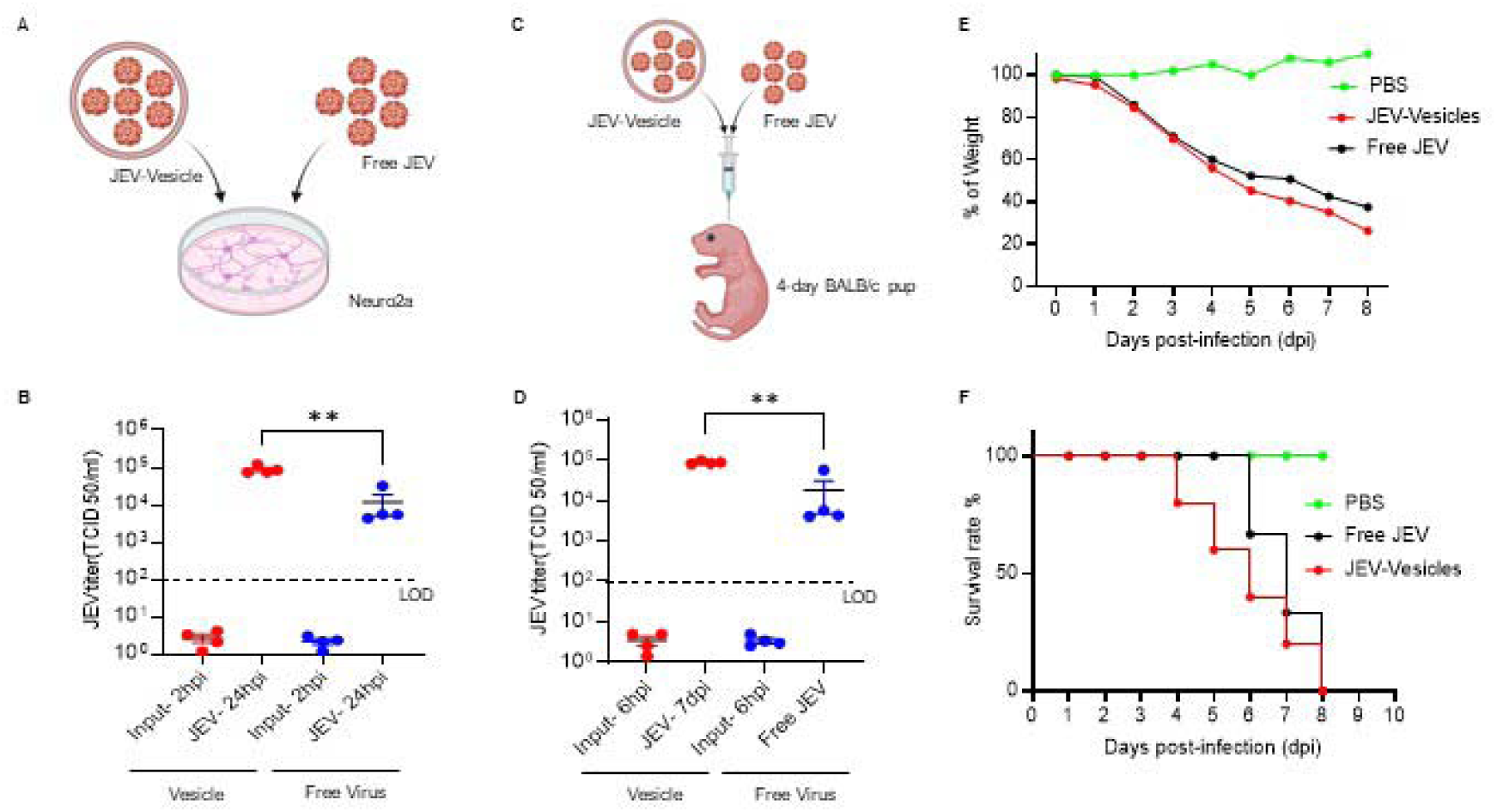
**JEV packaged within sEVs are infectious** (A-D) Equal amounts of JEV titer (based on TCID50/mL) from JEV-containing sEVs and free virus (released by lysis of vesicles using a non-detergent buffer) were used to infect (A, B) Neuro2a cell cultures and (C, D) 4-day-old BALB/c mouse pups via intracranial injection. (B) Neuro2a cells from each group were harvested at indicated time points, and viral titers were quantified by TCID50/mL. (D) Brain tissues from injected pups were collected and analyzed for viral titers. LOD = Limit of Detection for TCID50/mL assay. (E) % of weight loss and (F) Survival rate (%) were compared between groups receiving sEV-JEV versus free virus. Statistical significance between groups was assessed using unpaired t-tests; **pL<L0.01.

### 2.4. JEV-Induced sEV Release Occurs via a Ceramide-Dependent, ESCRT-Independent Pathway

Sphingomyelin (SM), a key membrane lipid enriched in neuronal cells, is enzymatically converted to ceramide by neutral sphingomyelinase 2 (nSMase2)—a well-characterized mechanism involved in the biogenesis of multivesicular bodies (MVBs) and small extracellular vesicles (sEVs) (Schneider *et al*, 2019; Schöl *et al*, 2024; Tohumeken *et al*, 2023). Given the neurotropism of JEV, we investigated whether this ceramide-driven, ESCRT-independent pathway facilitates viral release via sEVs during infection (schematic, Fig. 4A).

**Figure 4:**
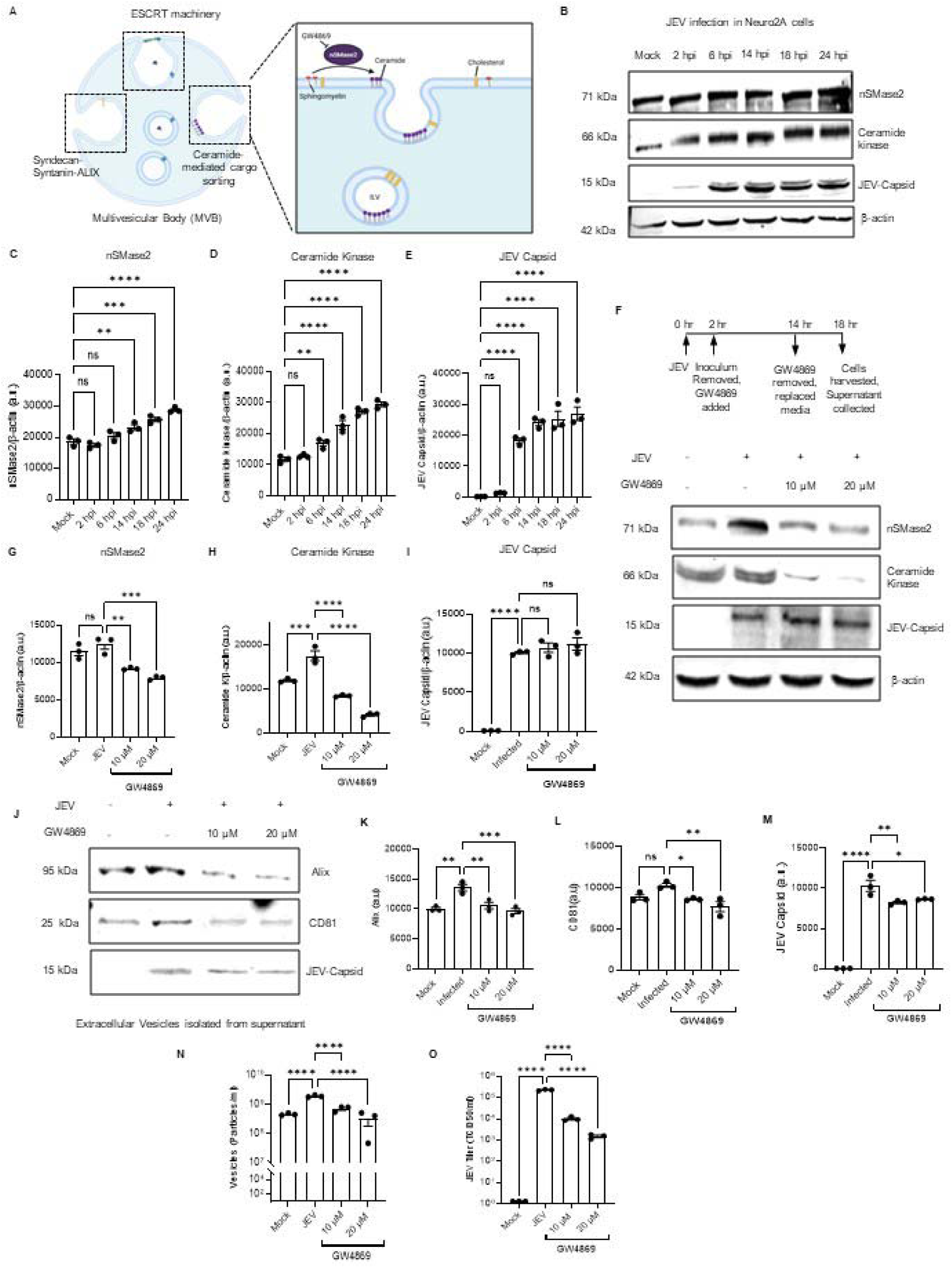
**JEV-Induced sEV Release Occurs via a Ceramide-Dependent, ESCRT-Independent Pathway** (A) Sphingomyelin is converted to ceramide by the enzyme nSMase2. Ceramide contributes to the formation of curvature on MVBs for intraluminal vesicle (ILV) formation. GW4869 inhibits nSMase2 activity. (B) Neuro2a cells inoculated with JEV show increased protein levels of nSMase2 and ceramide kinase, correlating with viral replication as indicated by JEV capsid expression in representative immunoblots. β-actin was used as a loading control. (C–E) Densitometric quantification of nSMase2, ceramide kinase, and JEV capsid protein levels from (B). (F) Immunoblots showing expression of the same proteins after treatment with GW4869 (10 µM and 20 µM for 12 hours) followed by harvesting cells infected with JEV at 18 hpi, and (G–I) corresponding densitometric analyses. From the same wells, supernatants were collected and sEVs were isolated. (J) Immunoblots showing expression of vesicle markers CD81 and Alix, along with JEV capsid protein in isolated sEVs, and (K–M) their densitometric quantification. An equal volume of samples was added to wells for normalization. (N) Vesicle concentration (particles/mL) was measured using nanoparticle tracking analysis, and (O) viral titer was determined from the same samples. Statistical analysis was performed using one-way ANOVA with multiple comparisons. *p<0.05, **pL<L0.01, ***pL<L0.001, ****pL<L0.0001, ns= not significant.

Immunoblot analysis revealed a time-dependent upregulation of nSMase2 and ceramide kinase expression in Neuro2a cells inoculated with JEV from 6 to 24 hpi, along with increased viral capsid expression (Fig. 4B–E). This suggest that JEV infection actively engages the ceramide pathway during replication. To functionally validate this mechanism, we employed GW4869—a pharmacological inhibitor of nSMase2—previously shown to block ceramide-dependent vesicle biogenesis (Choezom & Gross, 2022a). Treatment of Neuro2a cells with GW4869 at optimized doses (10–20 µM) and exposure duration (12 hours) significantly reduced the expression of nSMase2 and ceramide kinase, while not impairing JEV replication itself (Fig. 4F–I; Fig. S3A–F). The mRNA expression of nSMase2 was significantly reduced at both doses in the presence of the inhibitor (Fig. S3F). This indicates that ceramide pathway inhibition suppresses vesicle formation without cytotoxicity or interference with viral genome replication.

sEVs isolated from GW4869-treated cells displayed markedly reduced levels of canonical vesicle markers Alix and CD81, as well as JEV capsid protein (Fig. 4J–M). This was further supported by nanoparticle tracking analysis, which revealed a significant reduction in vesicle concentration (Fig. 4N; Fig. S3G-J), and by decreased JEV titer measured from the vesicle fractions (Fig. 4O). Collectively, these data demonstrate that JEV utilizes a ceramide-mediated, ESCRT-independent pathway for packaging and release via sEVs. Pharmacological blockade of this pathway significantly attenuates the release of virus-loaded vesicles without affecting intracellular replication, underscoring the specificity and functional importance of this egress mechanism.

### 2.5. JEV Infection Regulates Lipid Droplet Secretion to Facilitate Viral Egress via a Ceramide-Mediated Cargo Sorting Mechanism

Lipid droplets (LDs), which originate from the endoplasmic reticulum (ER), store neutral lipids and are increasingly recognized as key organelles in viral replication and assembly (Herker, 2024). Prior studies have demonstrated the involvement of LDs in JEV replication (Sarkar *et al*, 2021). In our study, confocal microscopy (Fig. 5A-C) and FACS (Fig. 5D) analysis revealed a dynamic modulation of LDs during JEV infection in Neuro2a cells. Specifically, both the number and size of LDs increased from 2 to 6 hours post-inoculation (hpi), followed by a progressive decline through 24 hpi (Fig. 5A–D, Fig. S4A).

**Figure 5:**
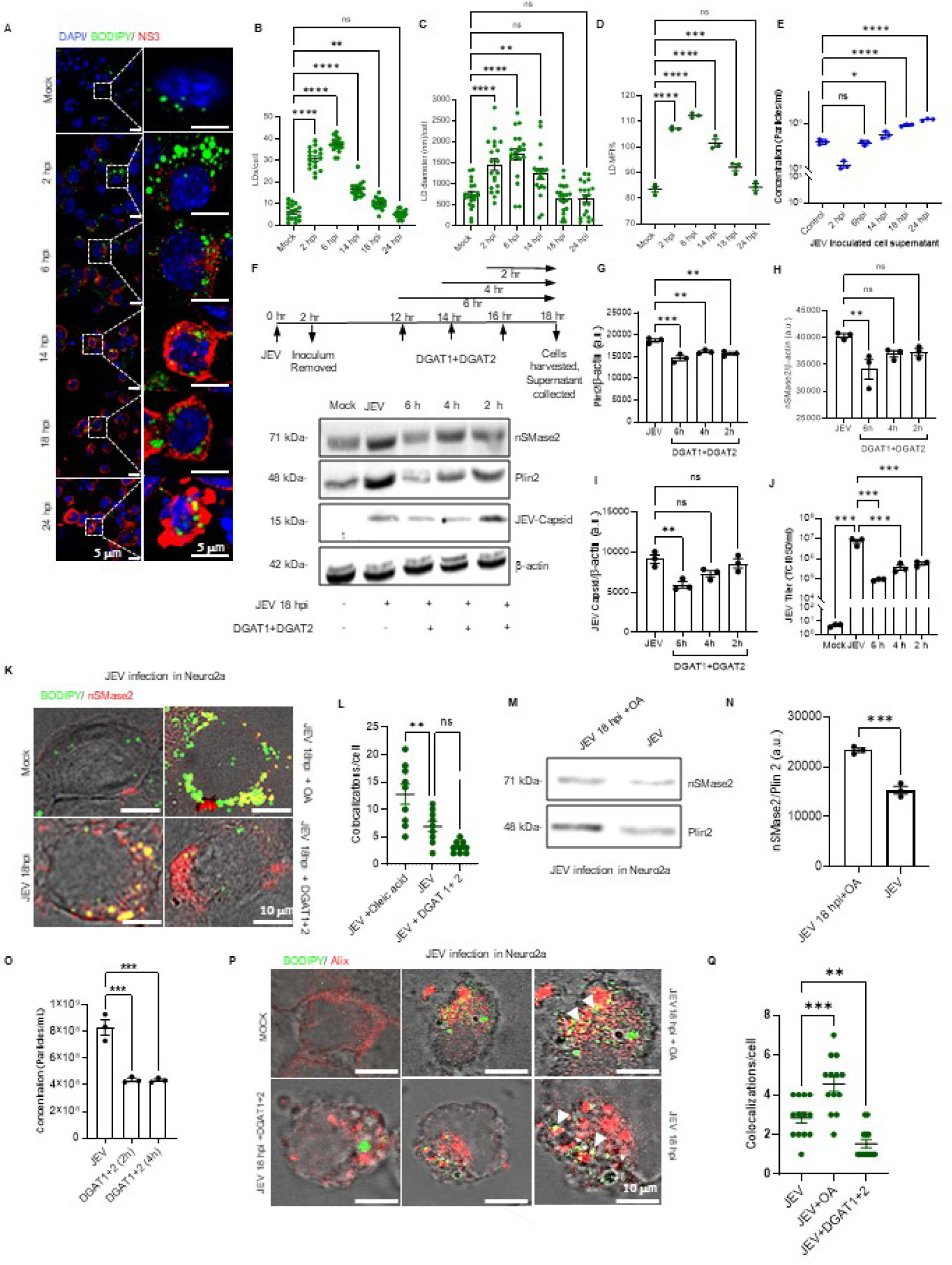
**JEV infection regulates lipid droplet (LD) secretion to facilitate viral egress via a ceramide-mediated cargo sorting mechanism.** (A) Representative confocal images showing LDs stained with BODIPY and viral replication marked by NS3 expression at 2, 6, 14, 18, and 24 hours post-inoculation (hpi). Enlarged views of representative cells are shown on the right. Scale Bar = 5Lµm. (B, C) Quantification of (B) the number of LDs per cell and (C) average LD diameter per cell across time points. (D) Flow cytometric analysis of BODIPY-stained LDs at indicated time points showing mean fluorescence intensity (MFI%). (E) Supernatants collected from the same wells were used to isolate sEVs, which were quantified using nanoparticle tracking analysis (NTA). (F) Neuro2a cells were treated with DGAT1 and DGAT2 inhibitors (LD biogenesis inhibitors) for 6-, 4-, and 2-hours during JEV infection and harvested at 18 hpi. Expression of nSMase2, Plin2, and JEV capsid protein was assessed by immunoblotting. β-actin served as the loading control. (G–I) Densitometric quantification of (G) Plin2, (H) nSMase2, and (I) JEV capsid protein from immunoblots shown in (F). (J) Infectious virus titers (TCID50/mL) were measured from sEVs isolated from the same treatment wells (K) Representative confocal images showing colocalization of BODIPY (LD marker) and nSMase2 staining under the following conditions: Mock, JEV (18 hpi) + Oleic Acid (OA), JEV infection (18 hpi), and JEV (18 hpi) + DGAT1+2 inhibitors (2 h). Scale Bar = 10Lµm. (L) Quantification of BODIPY and nSMase2 colocalization per cell. (M) LDs were isolated from JEV-infected (18 hpi) cells treated with or without OA and analyzed for nSMase2 expression by immunoblotting; Plin2 was used as the loading control. (N) Densitometric quantification of nSMase2 expression from (M). (O) NTA analysis of JEV-infected Neuro2a cells treated with DGAT1+2 inhibitor for different time points. (P) Representative confocal images showing colocalization of BODIPY (LD Marker), and Alix (MVB marker) staining under the following conditions: Mock, JEV (18 hpi) + OA, JEV infection (18 hpi), and JEV (18 hpi) + DGAT1+2-inhibitors (2 h). Scale Bar = 10Lµm (Q) The bar graph represents colocalization of BODIPY (LD marker), and Alix (MVB marker) at different time points in JEV-infected and DGAT1+2 treated Neuro2a cells. Statistical analysis was performed using one-way ANOVA with multiple comparisons and unpaired t-tests for pairwise comparisons. *p<0.05, **pL<L0.01, ***pL<L0.001, ****pL<L0.0001, ns= not significant.

Interestingly, the reduction in intracellular LD content beginning at 14 hpi coincided with a marked increase in sEV release, as quantified by nanoparticle tracking analysis (NTA) (Fig. 5E, Fig. S4B). This inverse correlation suggested a potential role for LDs in supplying membrane or cargo components for sEV-mediated viral egress. To further probe the functional role of LDs in this context, we inhibited LD biogenesis using inhibitors targeting DGAT1 and DGAT2, enzymes responsible for converting diacylglycerol (DAG) into triacylglycerol (TAG), the core component of LDs. Short-term treatments (2–6 hours) were optimized to minimize cytotoxicity and prevent disruption of JEV replication. Immunoblotting of treated cells showed decreased expression of Plin2, a structural LD membrane protein, confirming LD depletion (Fig. 5F–G). Notably, nSMase2 levels also declined under these conditions, along with viral titers in isolated sEVs; however, JEV capsid expression remained unaltered, signifying that the DGAT 1 & 2 dosage used never interfered with viral replication (Fig. 5H–J), suggesting a mechanistic link between LD abundance, ceramide-mediated sorting, and viral release. Confocal imaging of Neuro2a cells at 18 hpi showed colocalization of BODIPY-labeled LDs and nSMase2, which was enhanced upon Oleic Acid (OA) treatment—known to promote LD biogenesis—and diminished upon DGAT1+2 inhibition (Fig. 5K–L, Fig. S4C-D). This further supports the interplay between LD generation and ceramide pathway activation during JEV infection. To directly examine nSMase2 association with LDs, we isolated LD fractions from Neuro2a cells infected with JEV (18 hpi), with or without OA treatment. Immunoblotting revealed a significant enrichment of nSMase2 in the LD fraction upon OA stimulation, relative to Plin2 levels (Fig. M-N), indicating active recruitment of nSMase2 to LDs during infection.

To investigate whether sEV release is correlated with lipid droplet (LD) biogenesis, we quantified sEV concentration by NTA in JEV-infected Neuro2a cells treated with DGAT1+2 inhibitor for 2h and 4h (Fig. 5O, Fig. S5A-C). Our results revealed a marked reduction in sEV concentration in the presence of the DGAT1+2 inhibitor, suggesting a potential link between LD formation and sEV release. A recent study by Grenard *et al*. demonstrated a significant increase in CD63⁺ or Alix⁺ multivesicular bodies (MVBs) in irradiated pancreatic cancer cells, which also exhibited an increase in LDs following irradiation. These findings support a possible emerging role for lipid droplets in MVB formation (Grenard et al., 2021). Taking cues from this, we analyzed the mRNA expression and protein levels of the MVB marker Alix (Fig. S5D-F). Our data showed that both mRNA expression and protein levels of Alix were significantly elevated in JEV-infected samples over time compared to mock. Furthermore, confocal microscopy revealed that treatment with the DGAT1+2 inhibitor resulted in significant reduction in Alix levels at both 6- and 18-hours post-infection (Fig. 5P-Q, Fig. S5G-H). In addition, the degree of colocalization between LDs and Alix was notably reduced under DGAT1+2-inhibited conditions (Fig. 5Q). These observations suggest that LDs may colocalize with MVBs to provide essential lipid components necessary for MVB maturation. This process appears to be critical for efficient sEV formation and release. Together, these results demonstrate that JEV hijacks the host LD machinery to facilitate its egress via the ceramide-dependent, ESCRT-independent sEV pathway. The temporal regulation of LD dynamics appears to be closely coupled to the biogenesis and viral loading of sEVs during infection.

### 2.6. nSMase2 Regulates the Crosstalk Between Lipid Droplet Biogenesis and JEV Egress via Small Extracellular Vesicles

To further elucidate the role of nSMase2 in linking LD dynamics with JEV release via sEVs, both knockdown and overexpression experiments were performed in Neuro2a cells. Silencing of nSMase2 using specific shRNA resulted in significant reduction in nSMase2 protein levels compared to non-targeting (NT) controls, accompanied by a notable increase in Plin2 expression, indicating intracellular LD accumulation. However, no significant change in JEV capsid protein expression was observed (Fig. 6A–D). These findings suggest that while nSMase2 is dispensable for viral replication, it plays a critical role in mediating JEV egress.

**Figure 6:**
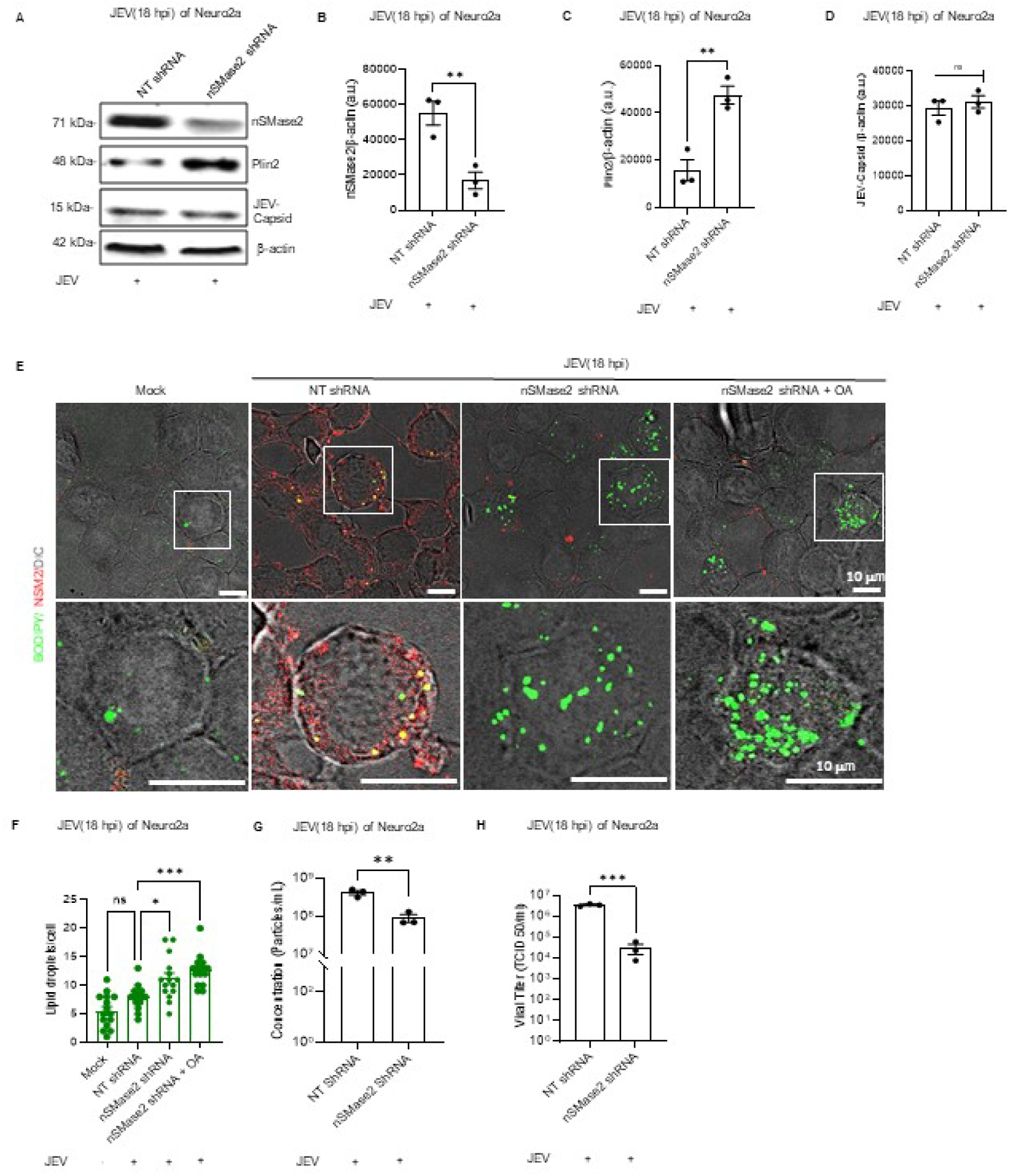
**nSMase2 knockdown inhibits JEV release via sEVs and enhances lipid droplet (LD) secretion.** (A) Representative immunoblot showing the expression levels of nSMase2, Plin2, and JEV capsid protein in Neuro2a cells transfected with non-targeting (NT) shRNA or nSMase2 shRNA, followed by JEV infection (18 hpi). β-Actin was used as the loading control. (B–D) Densitometric quantification of (B) nSMase2, (C) Plin2, and (D) JEV capsid protein, normalized to β-actin. (E) Representative confocal images showing colocalization of BODIPY (LD marker) and nSMase2 under the following conditions: Mock, JEV infection (18 hpi) + NT shRNA, JEV infection (18 hpi) + nSMase2 shRNA, and JEV infection (18 hpi) + nSMase2 shRNA + OA. Scale Bar = 10Lµm. (F) Quantification of LDs per cell corresponding to the images in (E). (G) Vesicle concentration (particles/mL) measured by nanoparticle tracking analysis (NTA) from the supernatant of same treatment groups. (H) Infectious JEV titer (TCID50/mL) measured from sEVs isolated from the same experimental conditions. Statistical analysis was performed using one-way ANOVA with multiple comparisons and unpaired t-tests for pairwise comparisons. *p<0.05, **pL<L0.01, ***pL<L0.001, ns= not significant.

Confocal microscopy revealed increased LD abundance and a more cytoplasmic localization pattern in nSMase2-depleted cells, as visualized by BODIPY staining (Fig. 6E–F). Notably, this was observed both with and without Oleic Acid (OA) treatment. In contrast, NT control cells infected with JEV displayed more peripheral colocalization of LDs with nSMase2 (as seen in Fig. 5K-L, Fig. 6E-F, Fig. S6A). NTA further revealed a significant decrease in sEV concentration in the nSMase2 knockdown group (Fig. 6G, Fig. S6B), and viral titers recovered from these vesicles were substantially reduced (Fig. 6H), reinforcing the role of nSMase2 in sEV-mediated viral export.

In contrast, overexpression of nSMase2 through pCMV6-nSMase2-GFP transfection led to elevated nSMase2 levels and a concurrent reduction in Plin2 expression, without affecting JEV capsid levels (Fig. 7A–D). This phenotype suggests that the enhanced turnover or utilization of LDs occurs in the presence of excess nSMase2. Confocal imaging showed reduced LD accumulation and increased membrane-associated nSMase2 in JEV-infected cells (Fig. 7E–F, Fig. S6C). Interestingly, upon OA treatment, a shift in nSMase2 localization from the plasma membrane to the cytoplasm was observed, accompanied by increased colocalization with LDs, suggesting redistribution in response to lipid enrichment.

**Figure 7:**
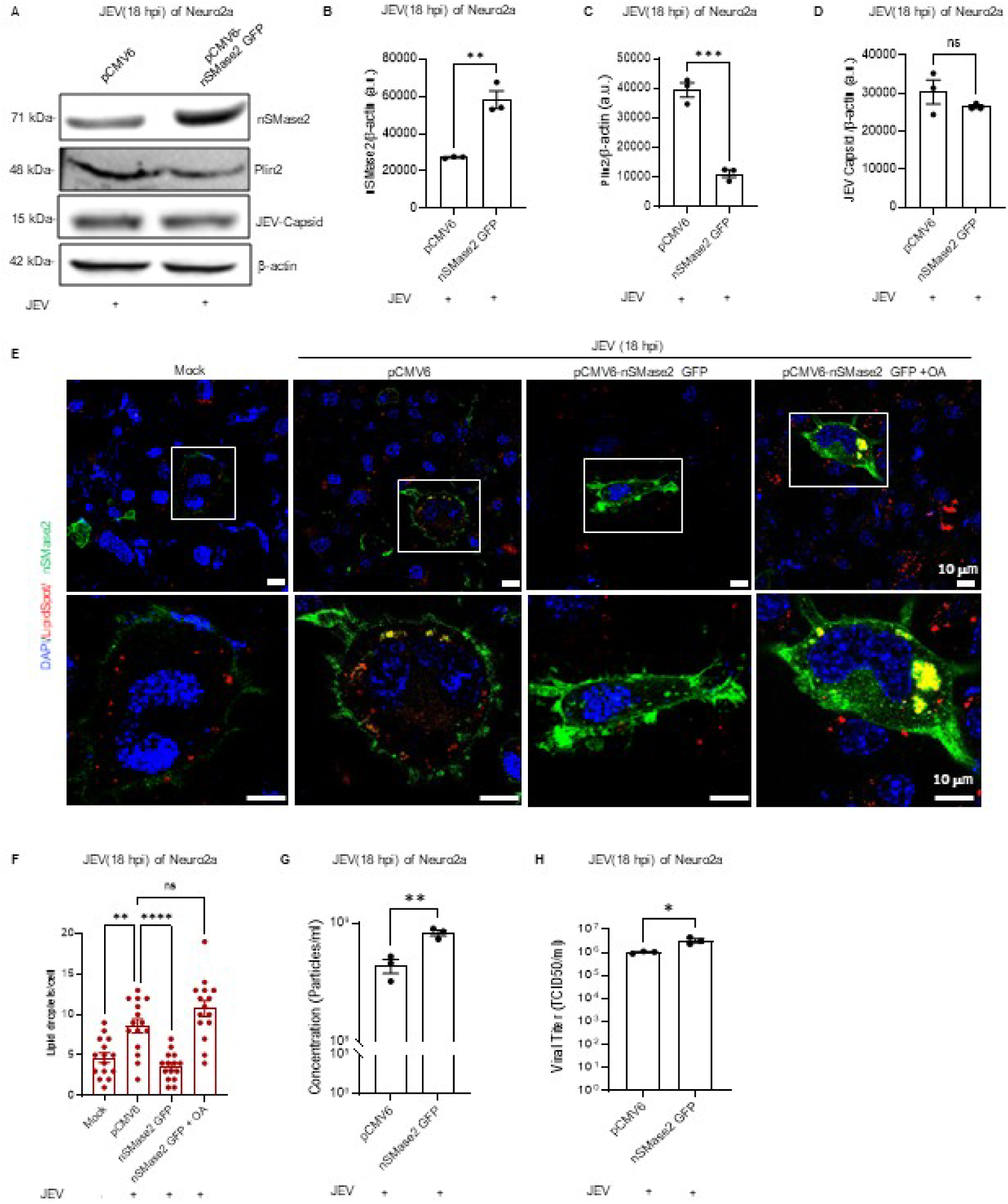
**Overexpression of nSMase2 suppresses LD secretion during JEV infection in Neuro2a cells.** (A) Immunoblot analysis showing protein levels of nSMase2, Plin2, and JEV capsid in Neuro2a cells transfected with either control vector (pCMV6) or nSMase2-GFP construct (pCMV6-nSMase2-GFP), followed by JEV inoculation (18 hpi). β-Actin served as the loading control. (B–D) Quantitative densitometry of (B) nSMase2, (C) Plin2, and (D) JEV capsid levels normalized to β-actin. (E) Confocal microscopy images illustrating the distribution of lipid droplets (LipidSpot staining) and nSMase2 under various conditions: Mock, JEV + pCMV6, JEV + nSMase2-GFP, and JEV + nSMase2-GFP + OA. Scale bar = 10Lµm. (F) LD count per cell corresponding to the imaging data shown in (E). (G) Quantification of sEVs (particles/mL) from the culture supernatant using nanoparticle tracking analysis (NTA). (H) JEV infectivity measured as TCID50/mL from isolated sEVs across experimental groups. Statistical analysis was performed using one-way ANOVA followed by multiple comparisons or unpaired t-tests, as appropriate. *pL<L0.05, **pL<L0.01, ***pL<L0.001, ****pL<L0.0001; ns = not significant.S

Consistent with these findings, sEVs isolated from nSMase2-overexpressing cells displayed a significant increase in vesicle concentration (Fig. 7G, Fig. S6D) and JEV titers (Fig. 7H), confirming enhanced vesicular export of infectious particles. Together, these gain and loss-of-function studies strongly support that nSMase2 acts as a central regulator of the ceramide-mediated, ESCRT-independent sEV biogenesis pathway exploited by JEV. Through modulating LD availability and localization, nSMase2 facilitates the incorporation of viral cargo into sEVs, thereby promoting efficient, non-lytic viral egress.

## 3 Discussion

Lipid droplets (LDs) and extracellular vesicles (EVs), which are distinct cellular components, have recently been shown to have overlapping functional landscapes. Traditionally regarded as separate—LDs serving as lipid storage depots and EVs as mediators of intercellular communication—both have now been implicated in key cellular and pathogenic processes (Amarasinghe *et al*, 2023; Genard *et al*, 2024). Emerging studies indicate that LDs and EVs play a crucial role in viral replication, intracellular trafficking, virion assembly, and unconventional secretion mechanisms (Sarkar *et al*, 2021; Herker, 2024; Zhang *et al*, 2017; Herrera-Moro Huitron *et al*, 2023). A significant conceptual advance emerged from the realization that LDs are not merely for lipid homeostasis and lipid storage sites, but also actively engage in membrane trafficking and organelle crosstalk (Menon *et al*, 2023; Fan & Tan, 2024). This includes interactions with endolysosomal compartments and multivesicular bodies (MVBs), the precursors of small EVs (sEVs), as shown by Genard et al. 2024, who provided the first direct experimental link between LD biogenesis and sEV release, mediated through RAB5C. In the context of viral infections, while both enhanced LD formation and sEV-mediated viral release have been observed independently, the molecular connection between these organelles during viral infection remains poorly defined. Our study addresses this gap by dissecting the mechanistic interplay between LDs and sEVs in Japanese Encephalitis Virus (JEV)-infected neuronal cells (Xia *et al*, 2023; Safadi *et al*, 2023; Miyanari *et al*, 2007; Giannessi *et al*, 2020).

Xiong et al. (2025) in their recent report showed that Japanese Encephalitis Virus (JEV) can be released non-lytically within small extracellular vesicles (sEVs) from PIEC, HeLa, and BHK-21 cells (Xiong *et al*, 2025). In line with their findings, our study provides additional evidence that JEV is similarly released non-lytically from neuronal cells, including Neuro2a, SHSY-5Y (a neuronal lineage), N9 (microglial cells), and primary cortical neurons, within vesicles ∼200 nm in diameter. Compared to JEV released through the secretory pathway, the JEV population packaged inside sEVs has a higher proportion of Premature Membrane protein (PrM) compared to Membrane protein, as indicated by a higher ratio of PrM/M. Vesicle-cloaked JEV were observed to be more infectious than the vesicle-free virions both *in vitro* and *in vivo* models. Collectively, these findings suggest that the sEV egress may serve as an adaptive mechanism, facilitating the egress of higher proportion of immature JEV particles and thereby conferring enhanced infectivity advantages. This process may enable a more rapid mode of viral transmission through vesicle-mediated mechanisms evading host antibodies.

Our data show JEV explicitly uses the ESCRT-independent pathway mediated by neutral sphingomyelinase 2 (nSMase2) and the ceramide biosynthesis pathway for its egress inside sEVs. The role of nSMase2 in sEV biogenesis is well-characterized in neurons, particularly in the release of tau protein and factors that promote neuronal survival and neuroprotection (Tallon *et al*, 2021a; Chen *et al*, 2024; Risner *et al*, 2023). Our findings reveal that JEV hijacks this endogenous machinery for its release via secreted extracellular vesicles (sEVs). Notably, although Xiong et al. reported that GW4869—a selective inhibitor of nSMase2—disrupted JEV replication and sEV release when used in pre-treatment, our study adopted a more targeted approach. We pre-treated infected cells with 10 µM and 20 µM of GW4869 for a minimum of 12 hours, ensuring cytotoxicity was avoided. Under these conditions, JEV replication remained unaltered, but its release in sEVs was substantially inhibited, underscoring the role of the ceramide-nSMase2 axis specifically in viral packaging and egress. The role of LDs in JEV replication and assembly has been previously reported (Sarkar *et al*, 2021; Ishida *et al*, 2019). We also observed a sharp increase in LD accumulation up to 6 hours post-infection (hpi), followed by a decline from 14 hours post-infection (hpi) onward. Interestingly, this temporal pattern inversely correlated with a marked increase in sEV release beginning around 14 hpi. These sequential dynamics prompted us to explore a potential functional linkage between LD dynamics and sEV-mediated viral release in JEV infection. This novel aspect may also extend to other viral systems.

It was a compelling observation that the reduction in LD number and size during JEV infection could be attributed either to a halt in LD biogenesis or to their functional consumption—(i) in supporting viral replication or (ii) in mediating sEV egress, or possibly both. Given the critical role of LDs in JEV replication, as highlighted in earlier studies, we adopted a cautious approach when inhibiting LD synthesis to ensure that viral replication remained unperturbed. Indeed, a short 2-hour treatment with dual DGAT1 and DGAT2 inhibitors—known to block LD biogenesis—did not affect JEV replication, yet significantly hampered the release of JEV within sEVs. Interestingly, at 18 hours post-infection (hpi), we observed that LDs were positioned peripherally and appeared to colocalize with nSMase2. This was further substantiated by our subcellular LD fractionation experiments, which revealed enrichment of nSMase2 in the LD fraction. In the context of metabolic disorders such as non-alcoholic fatty liver disease (NAFLD), often associated with type 2 diabetes and impaired immunity, LD accumulation is reported to correlate with ceramide (Cer) generation and nSMase2 expression on LD membranes (Schempp *et al*, 2024). However, in our model, we specifically detected this colocalization only at 18 hours post-infection (hpi). Since nSMase2 is known to traffic intracellularly between the Golgi and plasma membrane (El-Amouri *et al*, 2023), it may encounter LDs transiently during endosome maturation and multivesicular body (MVB) formation. LDs have previously been implicated in providing membrane lipids for MVB biogenesis, and given that nSMase2 is also crucial for intraluminal vesicle formation within MVBs, it is plausible that the observed colocalization at 18 hpi represents a functional convergence during sEV biogenesis. Direct physical or functional interactions between LDs and nSMase2 remain poorly understood. Our study, therefore, aimed to elucidate this interface in the context of JEV infection.

To dissect the functional relevance of nSMase2 in JEV trafficking, we employed shRNA-mediated knockdown of nSMase2 in infected cells. This resulted in a significant increase in the number of lipid droplets (LDs), as evidenced by enhanced expression of LD-resident membrane protein Plin2. This increase in LD biogenesis suggests a disruption in LD turnover and utilization. In control (non-target/scrambled) conditions at 18 hpi, LDs exhibited peripheral localization and showed spatial colocalization with nSMase2, possibly implicating their localization on the endosomal compartment (Choezom & Gross, 2022b; Genard *et al*, 2024). However, in nSMase2-depleted cells, LDs were redistributed across the cytoplasm and cell periphery without colocalization with nSMase2. Since nSMase2 catalyses the hydrolysis of sphingomyelin into ceramide, it plays a pivotal role in cellular processes. It is involved in the budding of intraluminal vesicles (ILVs) within multivesicular bodies (MVBs). Additionally, it contributes to small extracellular vesicle (sEV) biogenesis and cargo loading; its knockdown led to a substantial reduction in sEV secretion, as expected (Lee *et al*, 2024; Tallon *et al*, 2021a, 2021b). Notably, this reduction in sEV release was paralleled by diminished release of JEV, suggesting that nSMase2-driven ceramide enrichment is necessary for JEV export through the sEV pathway. This finding aligns with previous observations, where ceramide-mediated membrane curvature facilitates ILV formation—a prerequisite for exosome generation (Arya *et al*, 2022; Horbay *et al*, 2022; Wang *et al*, 2025).

The concurrent increase in LD numbers in nSMase2-deficient cells likely reflects a compensatory metabolic bottleneck. Under normal infection conditions, LDs may provide structural lipids, support membrane curvature, or serve as organelle contact surfaces that facilitate MVB maturation or cargo sorting. In the absence of efficient sEV formation, this downstream demand is attenuated, leading to an intracellular accumulation of LDs. Thus, our data suggest that LDs, although transiently associated with the egress process, are not directly involved in transporting JEV virions to the plasma membrane; instead, they serve in the assembly of MVBs. This conclusion is further substantiated by the inverse phenotype upon nSMase2 overexpression: a marked decrease in Plin2 levels and LD abundance, accompanied by enhanced sEV and viral release. We interpret this as LDs are actively consumed during MVB formation and viral packaging when the sEV biogenesis pathway is fully operational. It is also likely that the ceramide-enriched domains facilitated by nSMase2 may interact with LDs at membrane contact sites (MCSs), promoting lipid flux or localized membrane remodelling that drives MVB biogenesis.

In summary, our study investigates the poorly understood coordination between lipid droplets (LDs) and nSMase2 in sEV biogenesis and JEV egress. We found that LDs are not directly involved in viral egress but contribute to multivesicular body (MVB) formation. LDs and nSMase2 have independent yet synchronized roles in facilitating JEV trafficking and release. However, our reliance on *in vitro* models limits the understanding of LD-nSMase2 dynamics *in vivo*, especially within the brain’s complex environment. High-resolution and spatial transcriptomics, combined with organelle-specific analyses, could clarify the sequence of events during viral egress. Despite these limitations, our findings have translational relevance. The study suggests that organelle interactions are potential therapeutic targets, and studying EV formation and cargo could aid in the development of biomarkers for disease onset and severity. Future research using patient samples is crucial for translating these insights into detection and treatment strategies for JEV outbreaks.

## Author contributions

**Dr. Sourish Ghosh:** Conceptualization; project administration; investigation; formal analysis; methodology; resources; software; visualization; writing—original draft; writing—review and editing. **Bhaghyasree Mallick:** major data curation; formal analysis; investigation; methodology; writing—original draft; writing—review and editing. **Ankita Sarkar:** additional data curation; writing—original draft; editing original draft. **Ananya Mondal:** methodology; additional data curation; formal analysis. **Tamoghna Chakraborty:** additional data curation; formal analysis. **Khadija Khan:** data curation; formal analysis. **Dilip Kumar:** project management, review and editing. **Subhas Chandra Biswas:** project management, review and editing.

**Table 1:**
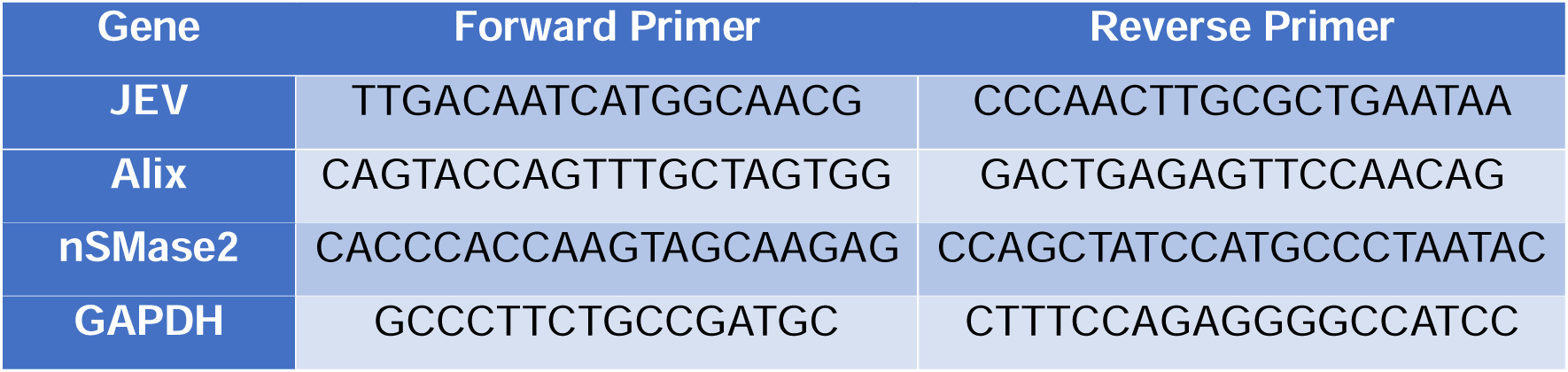
Primer list.

**Table 2:**
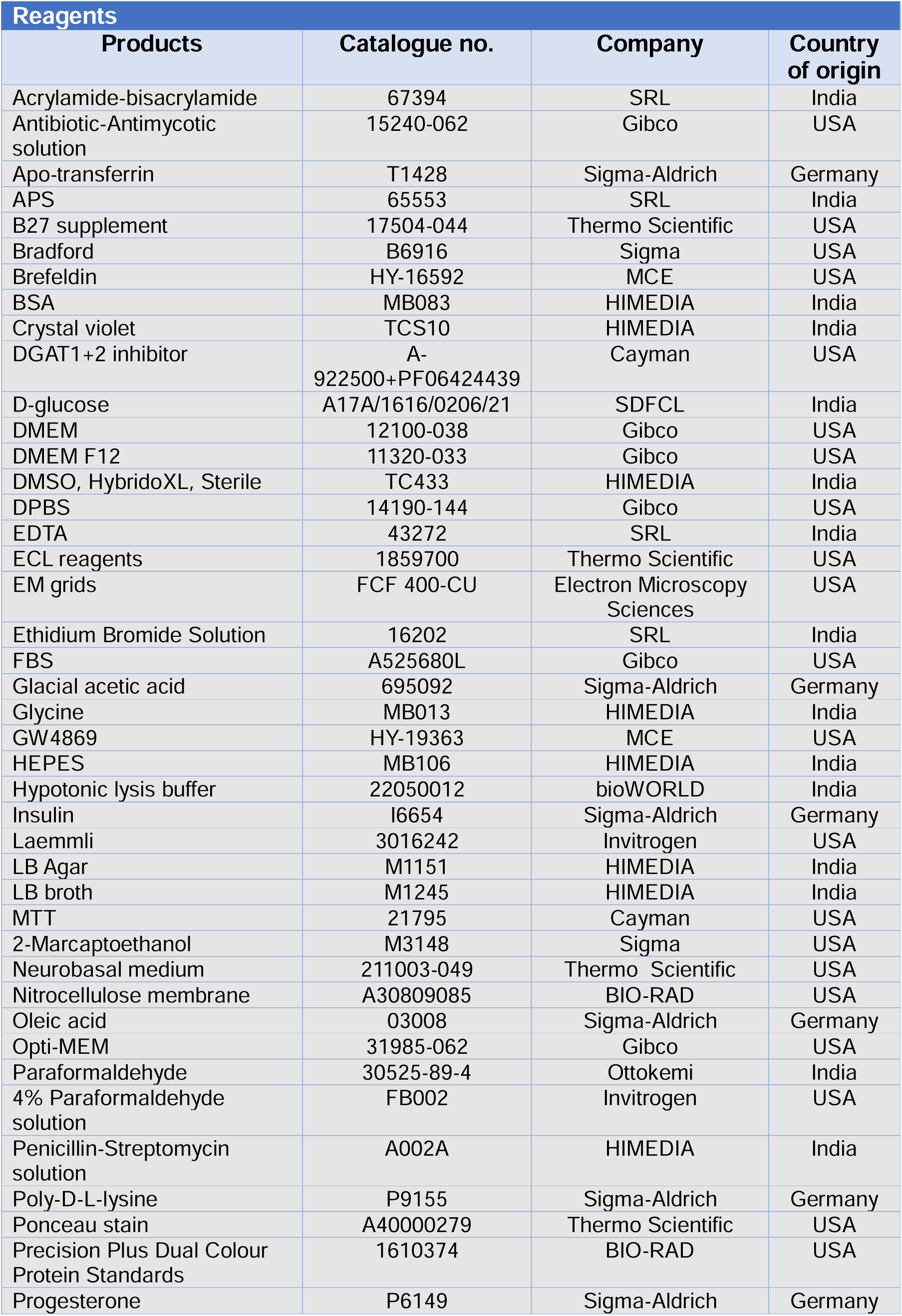
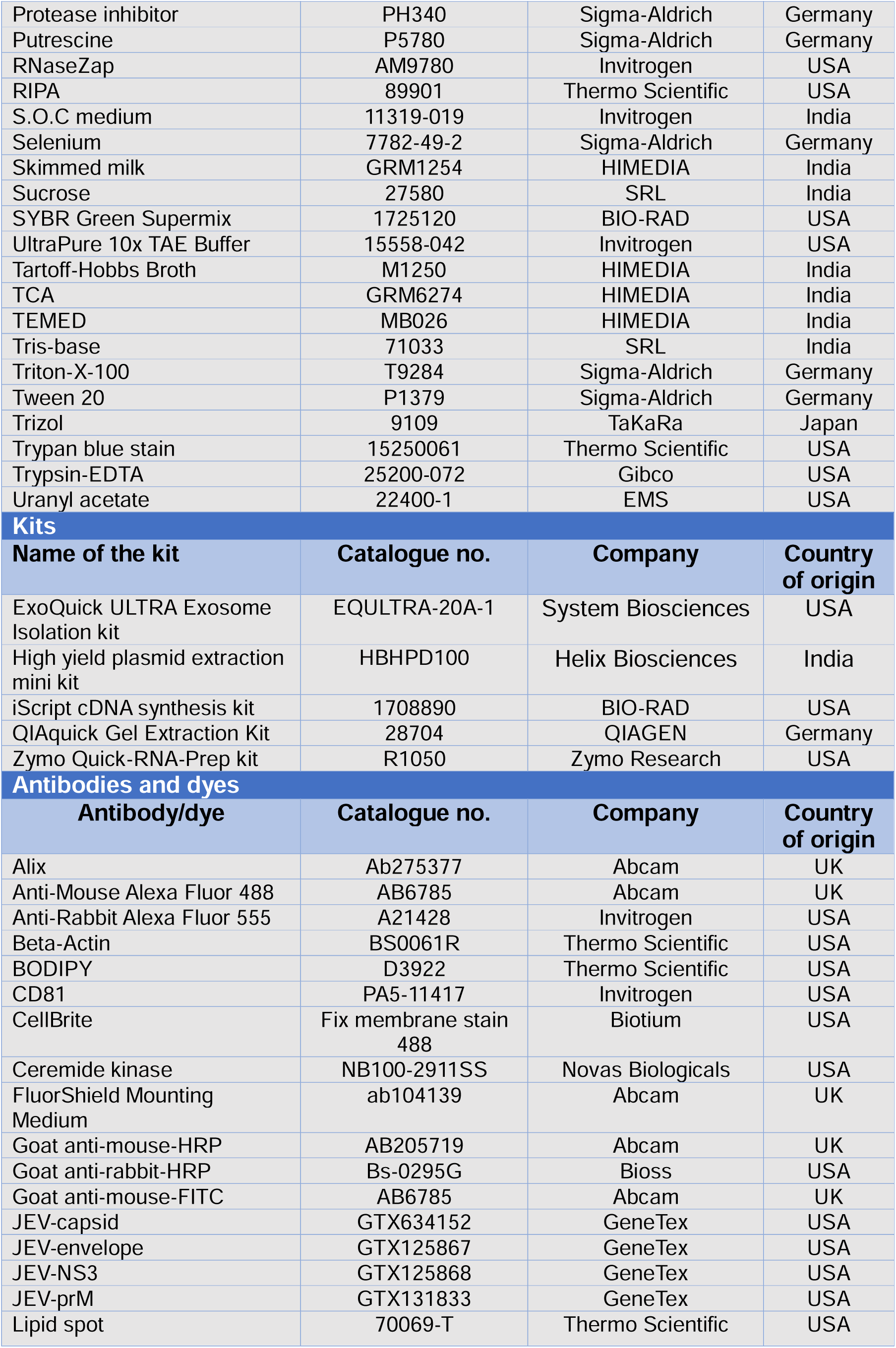
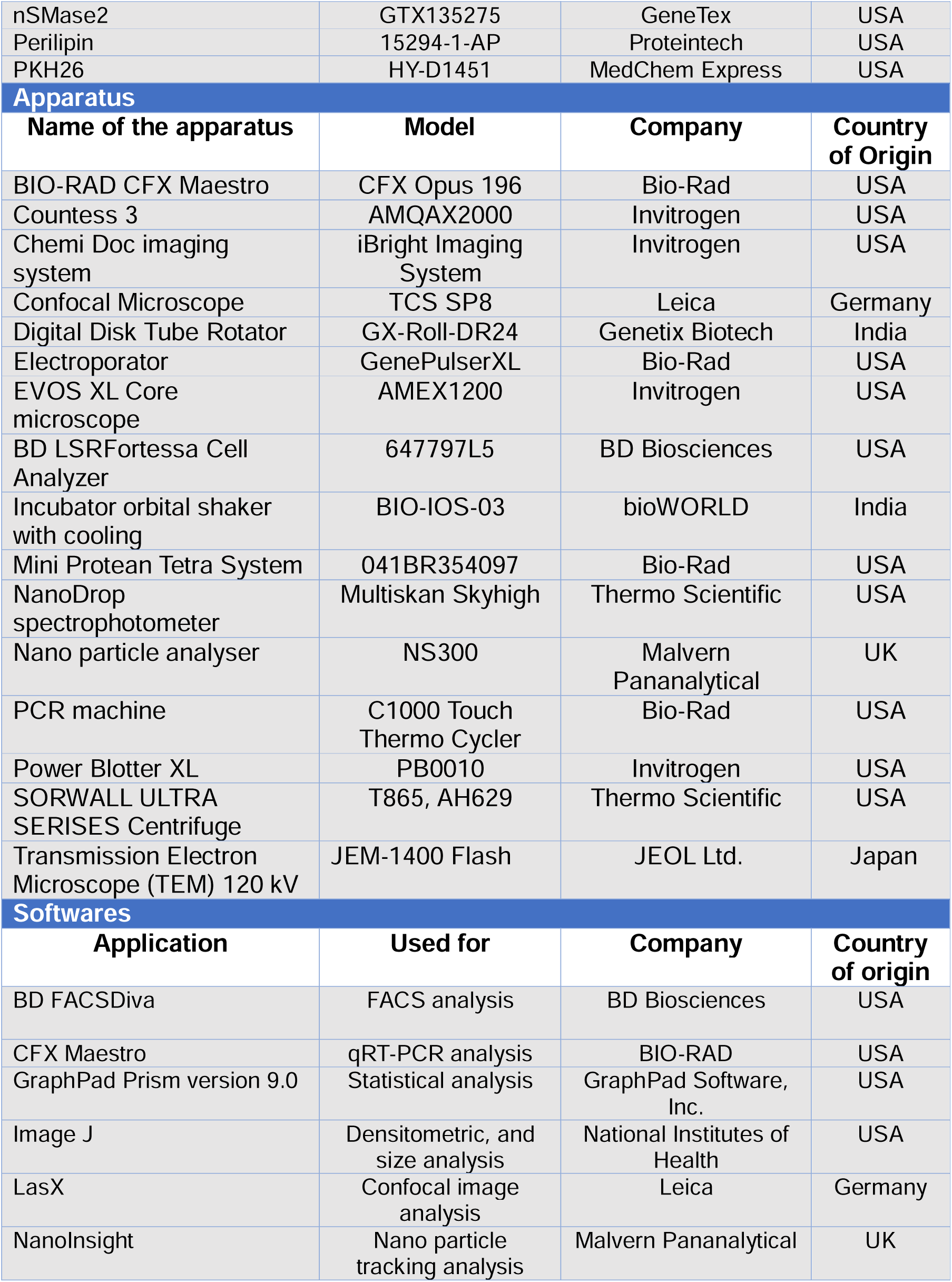
List of consumables, apparatus, and software

## Acknowledgement

This work was supported by CSIR-Focused Basic Research (FBR, No. – FBR070305) and the Department of Biotechnology (No. – BT/PR51484/MED/122/359/2024) awarded to SG. DK acknowledges the Annual Faculty Research Grant from Trivedi School of Biosciences, Ashoka University. AS is a DBT Research Associate (No. – DBT-RA/2023-24/Call-I/RA/07). BM, TC, and AM acknowledge the University Grants Commission (UGC) for their fellowships (JRF-NTA Ref no. 211610161998, 231620213546, 211610056569). KK acknowledges Ashoka University for the research fellowship. We also thank Mr. Sounak Bhattacharya, Senior Technical Officer, and Dr. Anirban Manna, Senior Technician at CSIR-Indian Institute of Chemical Biology (IICB), for their assistance with STED and Flow Cytometry, respectively. Electron Microscopy Facility of the Advanced Technology Platform Centre (ATPC), is managed by the Regional Centre for Biotechnology (RCB) and is funded by the Department of Biotechnology (Grant No. BT.MED-II/ATPC/BSC/01/2010). We also acknowledge that all the schematic figures in the manuscript were made using Biorender.com.

## Consent

All authors have agreed to the submission of this manuscript and confirm that it has not been published previously, nor is it under consideration for publication in any other journal.

## Data Availability statement

All data generated during the research work are included in the manuscript.

## Conflict of interest statement

The authors declare that they have no conflict of interest.

## Notes

### Competing Interest Statement

The authors have declared no competing interest.

